## Supplementary Material for "*O*-methylated N-glycans Distinguish Mosses from Vascular Plants"

**Citation:** Lastname, F.; Lastname, F.; Lastname, F. Title. *Biomolecules* **2021**, *11*, x. https://doi.org/10.3390/xxxxx

Academic Editor: Firstname Lastname

Received: date
Accepted: date
Published: date

**Publisher's Note:** MDPI stays neutral with regard to jurisdictional claims in published maps and institutional affiliations.

**Copyright:** © 2021 by the authors. Submitted for possible open access publication under the terms and conditions of the Creative Commons Attribution (CC BY) license (https://creativecommons.org/licenses/by/4.0/).

**Table S1**: **Identification of the 14 Da mass increment as methyl group rather than oxidation.** The measured and calculated mass increments are given. The 0.01 % (p = 0.0001) confidence interval was calculated from the eight values.

| Glycan | m/z | delta | | Statistics | |
|---|---|---|---|---|---|
| MMX | **1065.358** | | | average | 14.020 |
| | 1079.380 | 14.022 | | std. dev. | 0.008 |
| | 1093.390 | 14.010 | | conf. int 0.01 % | +/- 0.015 |
| MMXF | **1211.422** | | | | |
| | 1225.438 | 14.016 | | upper limit | 14.035 |
| | 1239.466 | 14.028 | | lower limit | 14.005 |
| MGnX | **1268.446** | | | of 0.01 % | |
| | 1282.463 | 14.017 | | conf.-interval | |
| | 1296.474 | 14.011 | | | |
| MGnXF | **1414.498** | | | calc. increment | |
| | 1428.519 | 14.021 | | methyl | 14.016 |
| | 1442.551 | 14.032 | | oxidation | 13.979 |

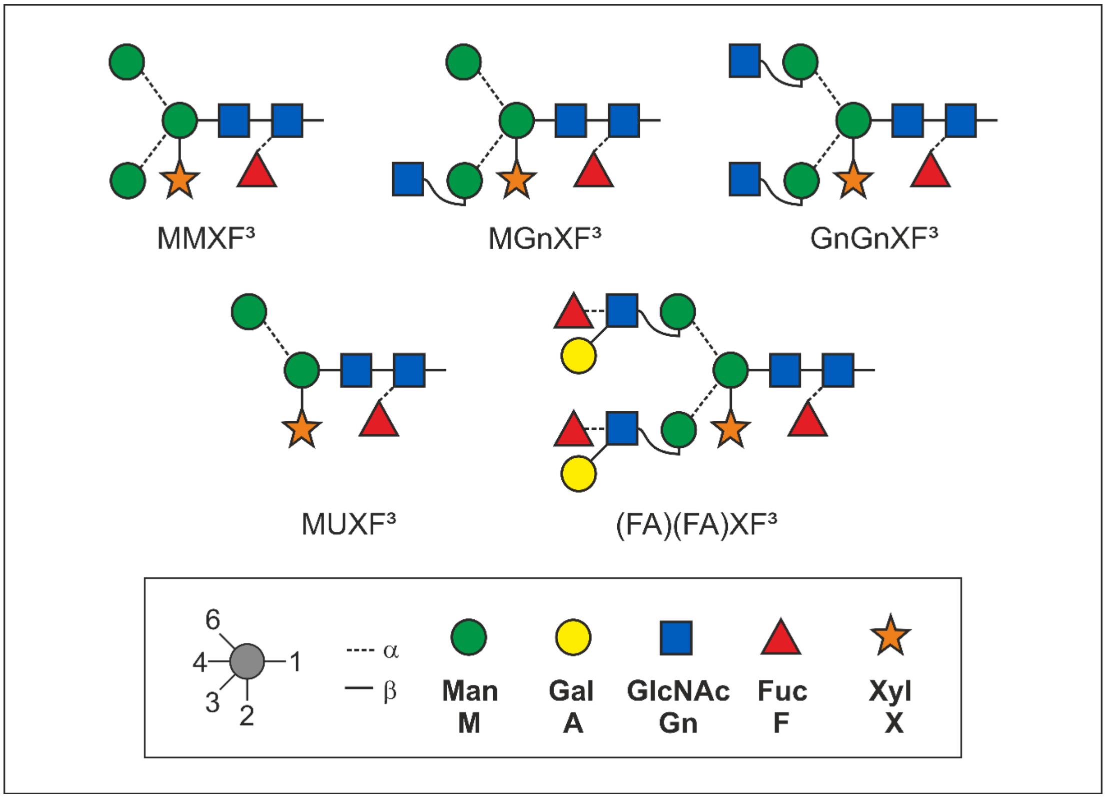


**Figure S1**: **Structures of plant N-glycans.** The proglycan nomenclature lists the terminal residues in a counter clockwise manner.

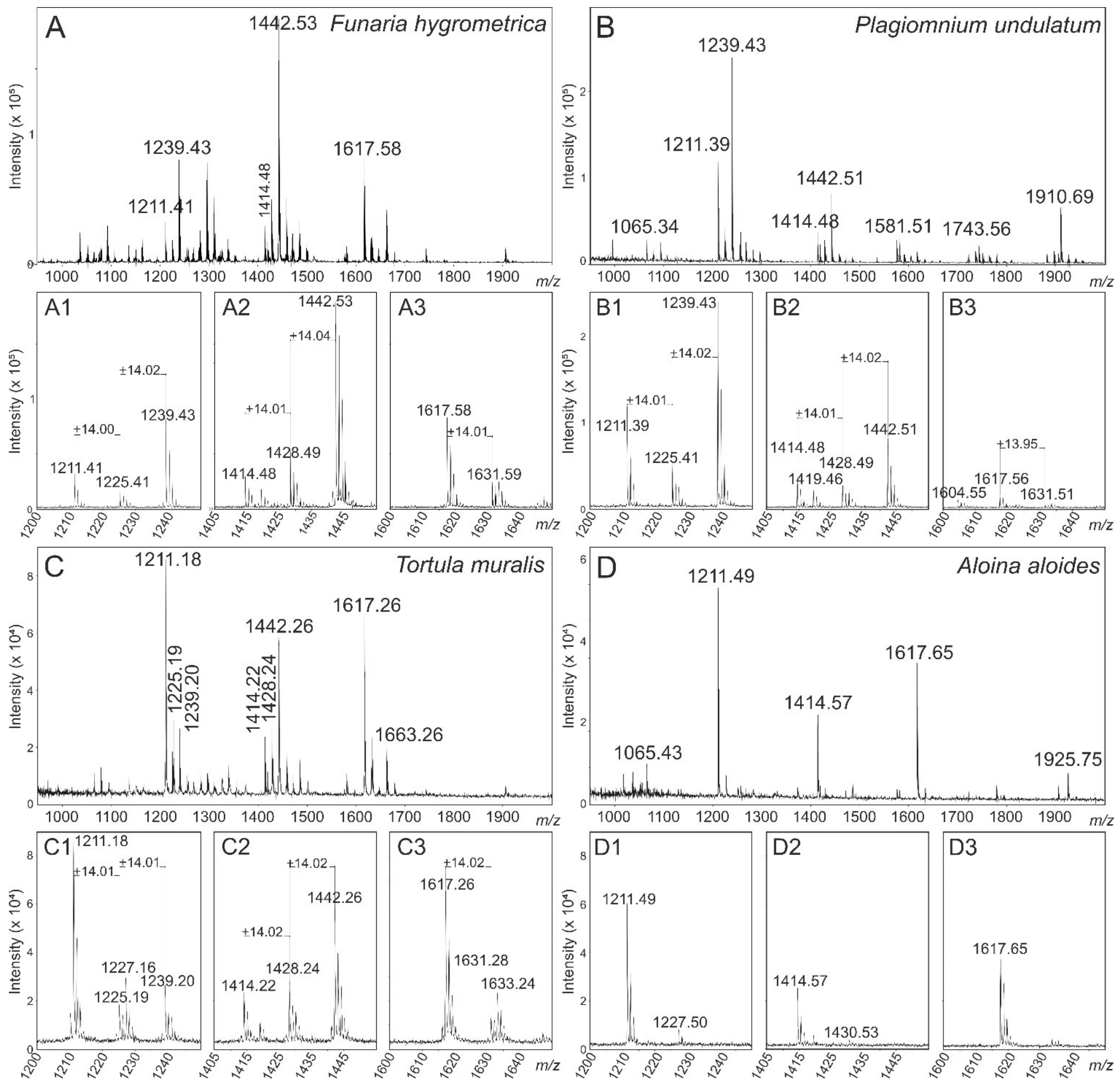


**Figure S2**: **MALDI-TOF measurements from different mosses.** MALDI-TOF measurements from native N-glycan pools from *Funaria hygrometrica* (**A**), *Plagiomnium undulatum* (**B**), *Tortula muralis* (**C**) and *Aloina aloides* (**D**). A1-D3 show zoomed in areas of the overall spectra.

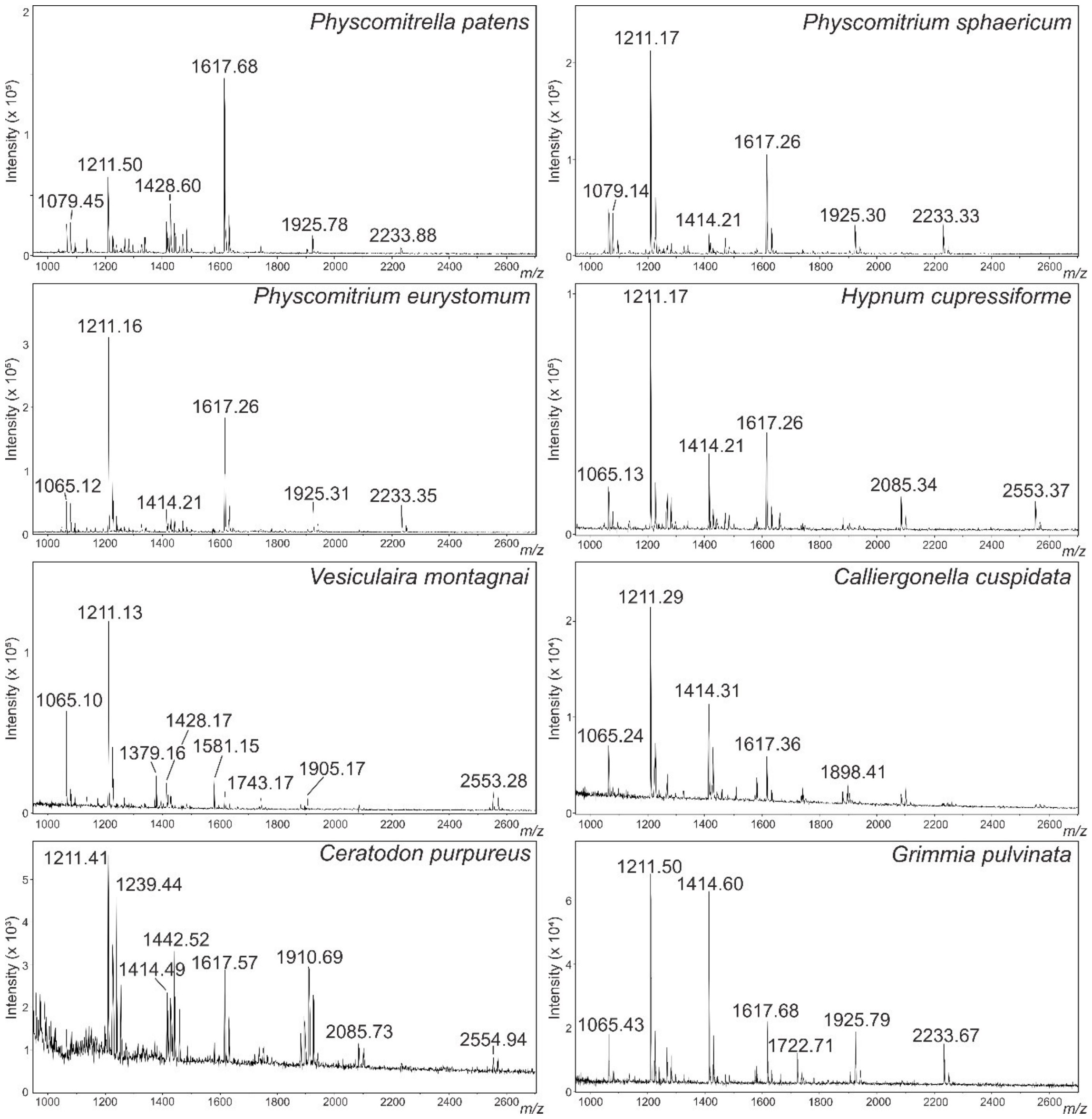


**Figure S3A**: **MALDI-TOF-MS of different moss species.** Whole glycan pools from the analysed moss species.

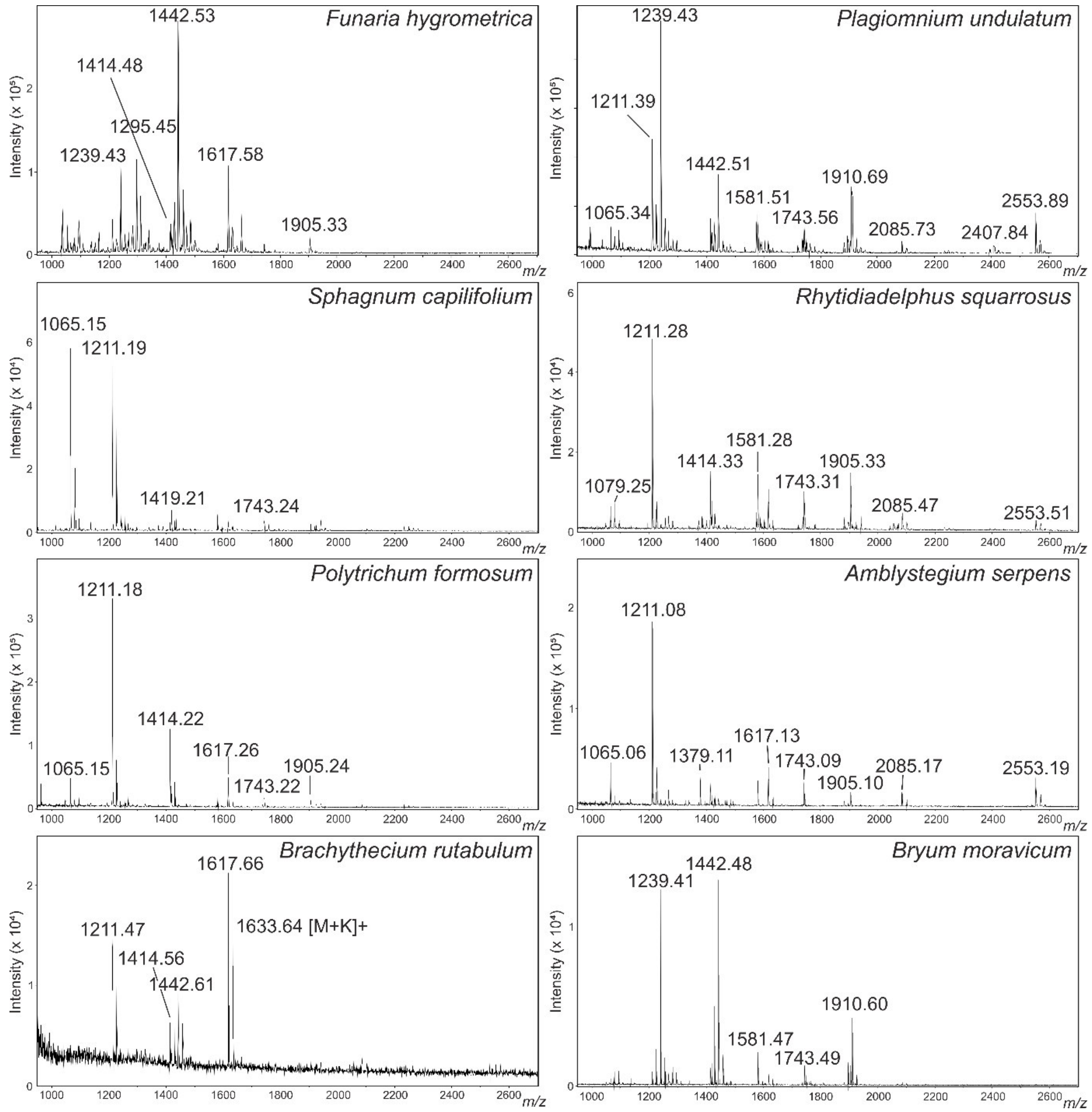


**Figure S3B**: **MALDI-TOF-MS of different moss species.** Whole glycan pools from the analysed moss species.

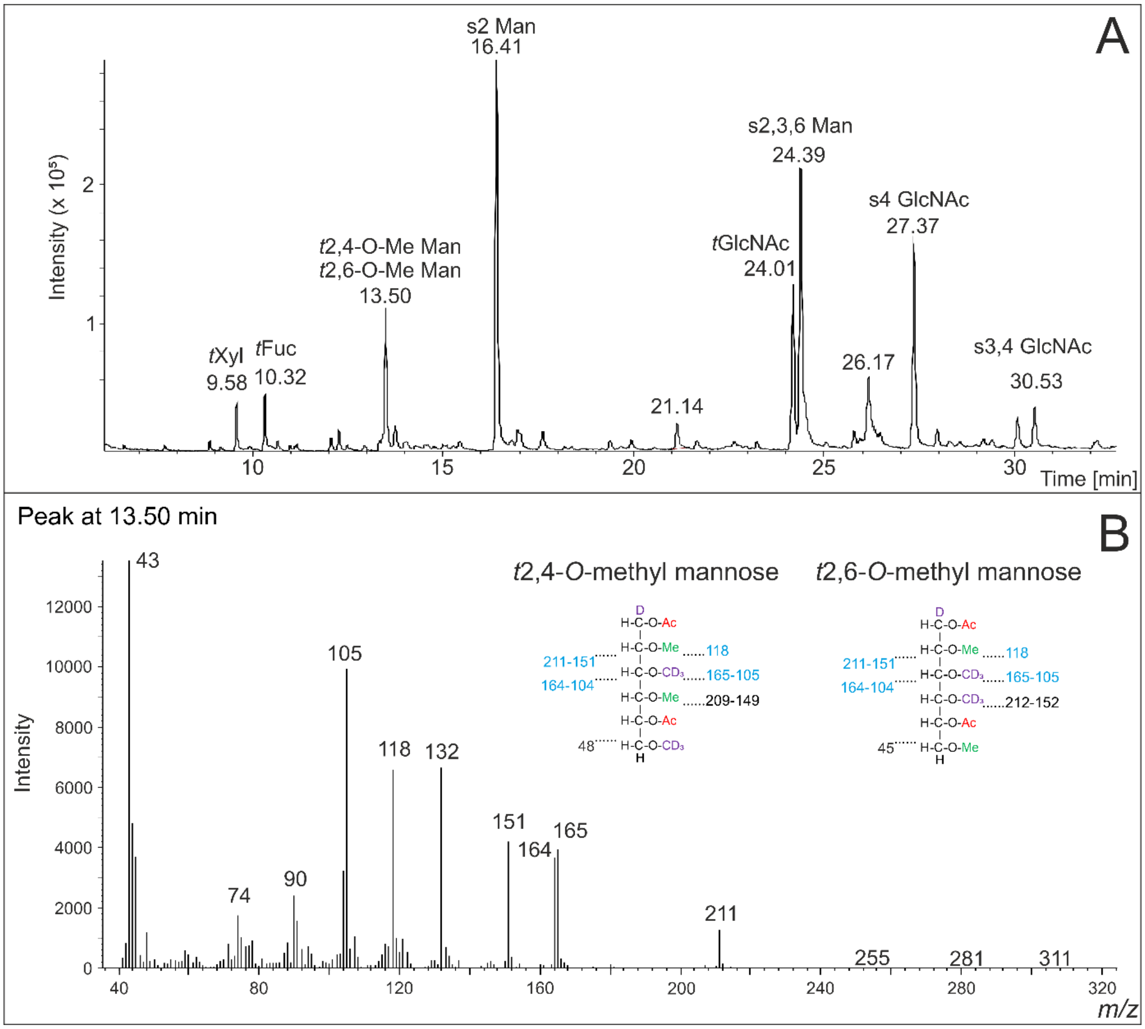


**Figure S4A:** (**A**): **GC-MS measurements of the HILIC-fractionated main glycan from *Funaria hygrometrica*.** The chromatogram from the first measurement of the sample treated with iodomethane-d3 revealed the doubly methylated mannose in a MGnXF. (**B**): The MS spectrum of this experiment could not clearly distinguish between 2,4-methylated mannose or a 2,6-methylated mannose.

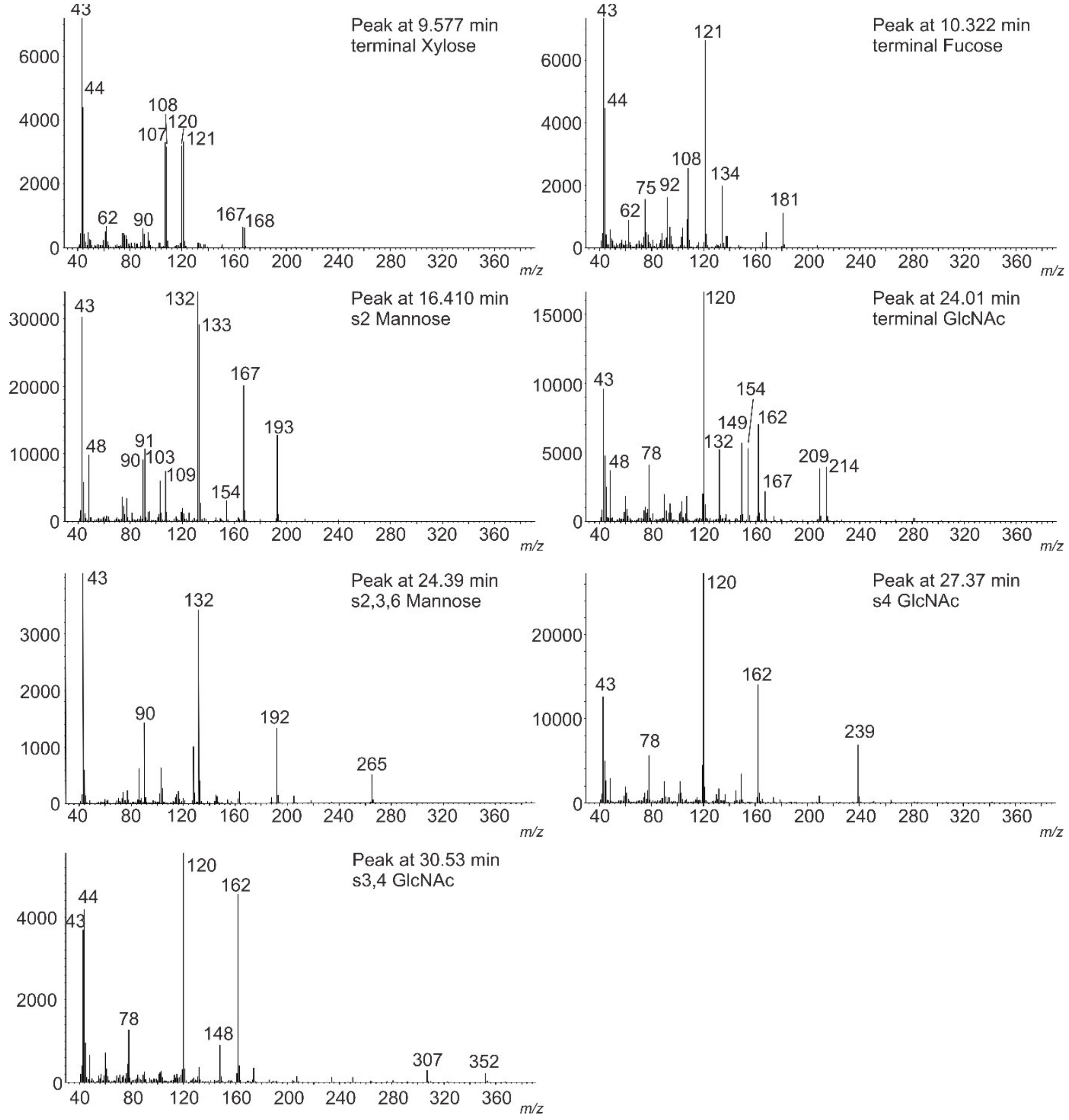


**Figure S4B**: **GC-MS measurements of the hydrolysed, peracetylated main glycan from *Funaria hygrometrica* treated with iodomethane-d3 before hydrolysis.** The resulting MS-spectra revealed the linkages between the monosaccharides as well as the possible methylation sites.

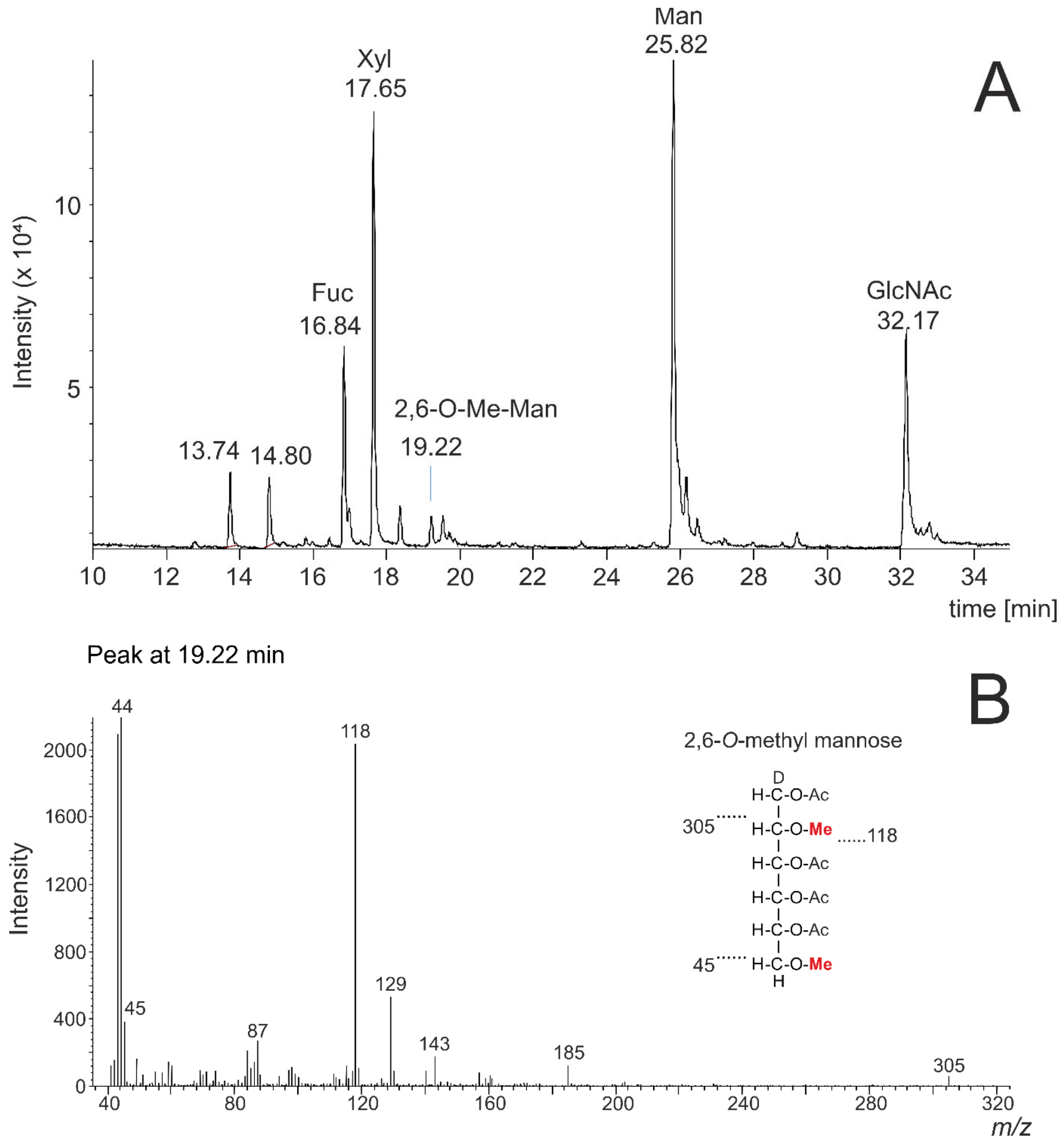


**Figure S5A**: **Localization of the methyl groups on the mannose of MGnXF of *Funaria hygrometrica*.** (**A**): GM-MS chromatogram of the hydrolysed and peracetylated main glycans from *Funaria hygrometrica* without the treatment of iodomethane-d3. (**B**): MS spectrum of the peak "19.22 min". With these measurements the methylation sites of the mannose were shown.

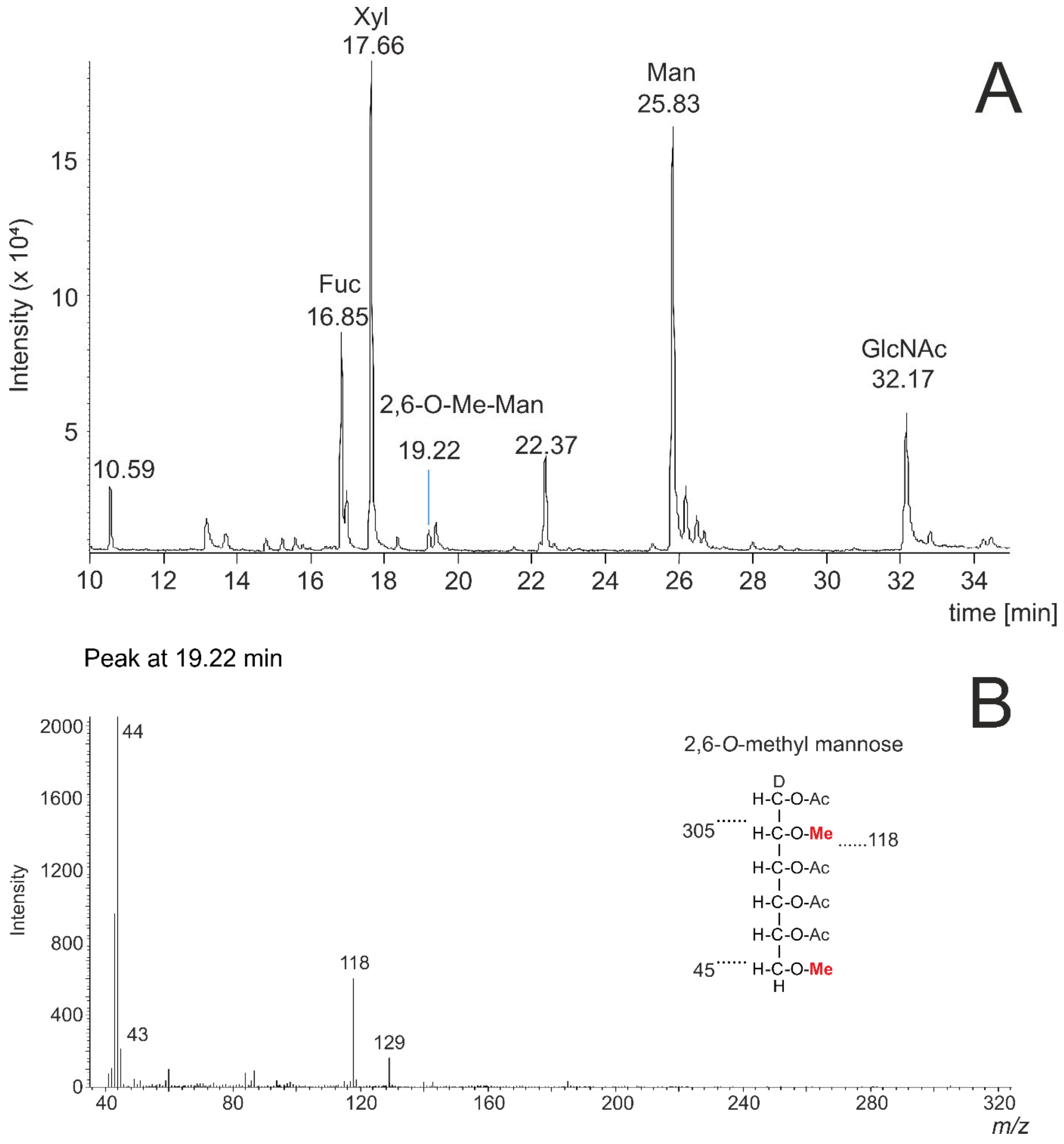


**Figure S5B**: **Localization of the methyl groups on the mannose of MMXF of *Plagiomnium undulatum*.** (**A**): GM-MS chromatogram of the hydrolyzed and peracetylated main glycans from *Plagiomnium undulatum* without the treatment of iodomethane-d3. (**B**): MS spectrum of the peak "19.22 min". With these measurements the methylation sites of the mannose were shown.

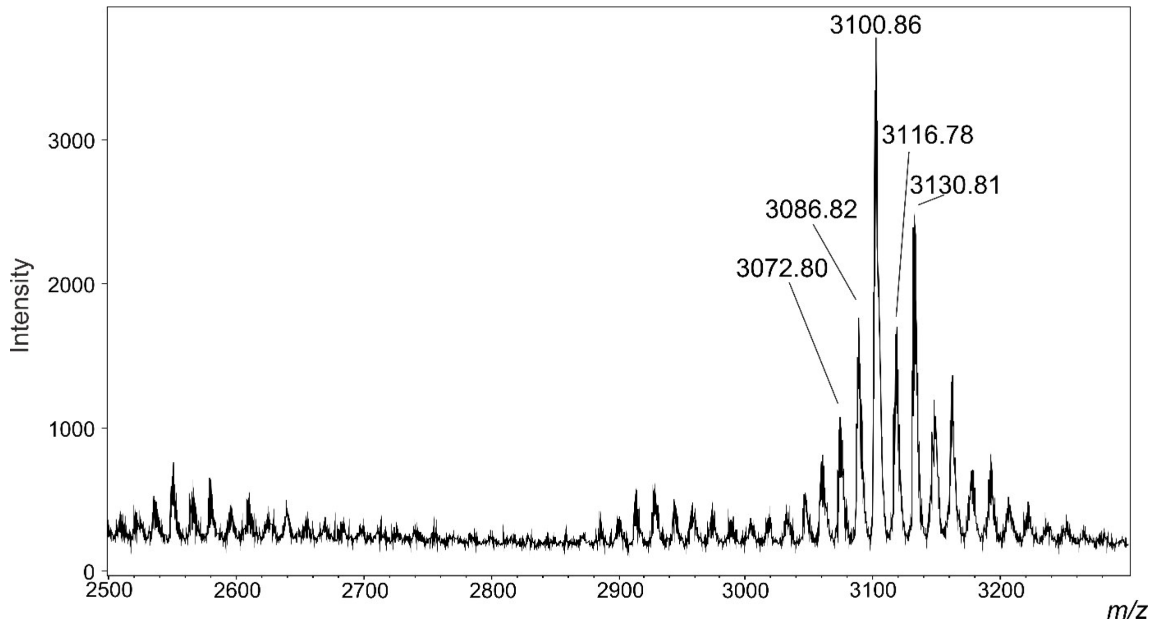


**Figure S6**: **MALDI-MS spectrum of permethylated 2553 [M+Na]$^+$ of *Plagiomnium undulatum*.** 3100.86 [M+Na]$^+$ represents the permethylated glycan.
